# Hebbian and homeostatic plasticity mechanisms are segregated in sub-types of layer 5 neuron in the visual cortex

**DOI:** 10.1101/2022.02.11.480060

**Authors:** Anurag Pandey, Neil Hardingham, Kevin Fox

**Affiliations:** Cardiff University

**Keywords:** CaMKII, TNFα, Intrinsic bursting, Regular spiking, Monocular deprivation

## Abstract

Cortical layer 5 contains two major types of projection neuron known as IB (intrinsic bursting) cells that project sub-cortically and RS (regular spiking) cells that project between cortical areas. We studied the plasticity properties of RS and IB cells in the visual cortex during the critical period for ocular dominance plasticity in mice. RS neurons exhibited synaptic depression in response to both dark exposure (DE) and monocular deprivation (MD), and their homeostatic recovery from depression was dependent on TNFα. In contrast, IB cells demonstrated opposite responses to DE and MD, potentiating to DE and depressing to MD. IB cells’ potentiation depended on CaMKII-autophosphorylation and not TNFα. IB cells showed mature synaptic properties at the start of the critical period while RS cells matured during the critical period. Together with observations in somatosensory cortex, these results suggest that differences in RS and IB plasticity mechanisms are a general cortical property.

**Significance Statement:** The neocortex contains cells that project to different locations in the brain. In this study we show that neurons projecting to different target locations exhibit different synaptic plasticity mechanisms. Cortically projecting cells show synaptic depression and homeostatic up-regulation, subcortically projecting cells show classical Hebbian potentiation. This is important because it implies that the way a cortical neuron responds to experience and encodes information depends on the neuronal subcircuits in which it is embedded. We show that ignoring this distinction leads to erroneous conclusions regarding plasticity time-course and significance. These findings constitute an important step toward understanding how learning and memory is organized within subcircuits in the cerebral cortex.

## Introduction

The cerebral cortex contains a diversity of excitatory neuronal subtypes. During development, a small set of relatively homogeneous progenitors give rise to several different types of adult neuron, expressing different molecules, displaying different dendritic morphologies, connecting to different neuronal circuits and projecting to different targets in the brain (1, 2). Among the projection cells, layer 5 neurons can be classified broadly into those that project sub-cortically, to targets such as the superior colliculus and pontine nuclei and those that project cortico-cortically between different cortical areas. The projections of these neurons correlate with their intrinsic membrane properties, their neuronal morphology and their intracortical connectivity (3–5). The sub-cortically projecting neurons produce bursts of action potentials when depolarised (intrinsic bursting, IB cells), have highly branched apical dendrites and receive input from several cortical columns, whereas the cortico-cortically projecting neurons tend to fire regular trains of action potentials when depolarised (regular spiking RS cells)(5, 6) have little or no branching of the apical dendrite and mainly receive input from within the cortical column (5, 7). In addition, RS cells either project to striatum and cortex or just cortico-cortically (8). Recently, studies in barrel cortex have shown that the differences exhibited by layer 5 RS and IB cells also extend to their plasticity mechanisms (9). While IB cells show αCaMKII-dependent potentiation and little depression during patterned whisker deprivation, RS cells show strong experience-dependent depression and little αCaMKII-dependent potentiation, instead exhibiting TNFα-dependent homeostatic plasticity, which is characteristic of synaptic scaling (9, 10).

These studies raise the question of whether the divergent plasticity mechanisms seen in layer 5 neurons are a specialised feature of barrel cortex or a broader property of cortical organisation. To test this idea we needed to study plasticity in RS and IB cells in at least one other cortical area. Given that the visual cortex is highly plastic and that experience-dependent plasticity has been studied extensively in this structure (11, 12), we sought to determine whether layer 5 neurons showed different plasticity mechanisms in the IB and RS cells of the visual cortex.

Plasticity in the visual cortex is thought to operate partly via synaptic scaling (13, 14), a process originally studied by application of TTX to neuronal cultures *in vitro* (13, 15). Two conditions are known to lead to TNFα-dependent up-scaling processes in the visual cortex; dark exposure *in vivo* causes up-scaling of synaptic weights (16) and eye-enucleation causes enlargement of the surviving dendritic spines on dendrites that have lost spines (17). However, other studies have implicated NMDA receptors and CaMKII auto-phosphorylation in plasticity induced by dark exposure, both of which would suggest an LTP-type mechanism is also involved (18–21). Both Hebbian and synaptic scaling mechanism have mainly been studied in layer 2/3 to date, so an additional question arises about which, if either, apply to layer 5.

We found that the mechanisms employed by neurons for TNFα-dependent homeostatic responses are specific to cortical projection subtypes. RS cells projecting cortico-cortically, do indeed exhibit homeostatic up-scaling dependent on TNFα processing following experience-dependent synaptic depression; however, IB cells projecting sub-cortically do not; instead, they show a response that depends on αCaMKII-autophosphorylation, a key molecular process required for LTP. The subtype divergence of plasticity mechanism is therefore not a peculiarity of barrel cortex but extends to visual cortex, strongly suggesting plasticity subtypes are a cortex-wide phenomenon.

## Results

### IB and RS Neuron Characteristics

IB and RS neurons were characterised electrophysiologically online from their firing patterns in response to somatic current injection (Fig. 1A). We found that the electrophysiological classification correlated well with several morphological characteristics. Sholl analysis revealed that dendrites were generally more branched in IB than RS neurons (F_(1,1)_=41.24, p<0.0001) and that this was true for basal, apical and apical oblique dendrites (Fig. 1B). In addition, the span of the apical tuft in layer I was consistently far broader in IB cells (average = 280μm) than in RS cells (average = 212μm) and significantly different (t_(16)_=1.78, p<0.05), which resulted in IB cells’ apical dendrites spanning a greater horizontal distance than their basal dendrites (80% of cases) while RS cells’ apical tufts spanned the same (within 10%) or a smaller radius than the basal dendrites (100% of cases). We also found that the electrophysiological identity of the neurons correlated with their projection targets (Extended Data Fig. 1). Neurones projecting to the superior colliculus showed a burst of spikes in response to somatic current injection (93%). Similarly, neurons that projected to the opposite hemisphere of the visual cortex had a 91% chance of exhibiting a regular spiking response to somatic current injection (Extended Data Fig. 1). These characteristics confirm that our electrophysiological categorisation of IB and RS neurons was consistent with those of previous studies (3-5, 22).

**Figure. 1.**
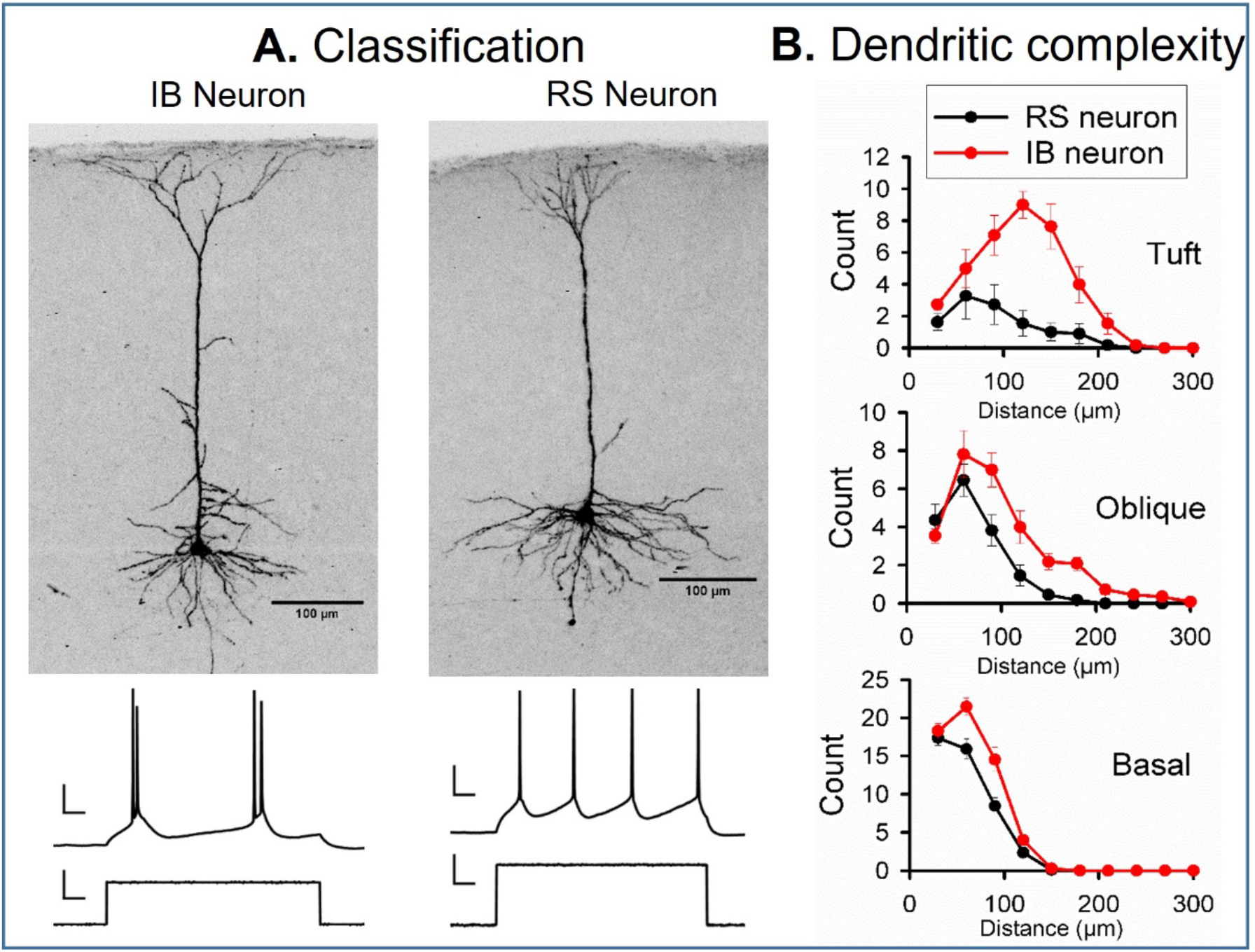
IB and RS neuronal properties. **A:** Example morphological differences in electrophysiologically identified IB and RS cells. The spike discharges (scale bars 20mV and 100ms) are produced from somatic current injection (scale bars 20pA and 100ms). **B:** Sholl plots for apical tuft, oblique and basal dendrites from RS (black) and IB cells (red). RS neurons have significantly fewer branches than IB cells at apical tuft (t_(20)_=5.03, p<0.0001) apical oblique (t_(20)_=2.87, p<0.01) and basal dendritic locations (t_(20)_=3.13, p<0.01).

We discovered one morphological feature concerning spine type that has not previously been reported to be different between RS and IB cells. The distribution of spine types was different in RS and IB cells (Extended Data Fig. 2). RS cells had a greater proportion of long thin spines (average 30% in RS cells vs 7% in IB), while IB cells had a greater proportion of mushroom shaped spines (average 64% in RS cells vs 83% in IB). These two spine types made up the majority of the total for both cell types, with stubby and filopodial spine types constituting only 6-10%. The difference in the proportion of spine types between RS and IB cells was highly statistically significant (χ^2^=23, df=2, p<0.001).

We also found a difference in spine density in IB versus RS cells. A 2-way ANOVA showed an effect of cell type on spine density (F_(1,1)_=18.12, p<0.0002) and an interaction between the location of the spines (apical versus basal) and the cell type (IB versus RS) (F_(1,1)_=5.46, p<0.03). In general, IB cells had a lower spine density (by 28%) than RS cells (Table 1), but this was most noticeable on the apical dendrites where the spine density (of 0.52 spines per micron) was approximately 40% lower than on IB cell basal dendrites or any location on RS cells (post-hoc t-test, α=0.05) (Extended Data Fig. 2). Taken together with the spine category differences, these observations suggests that RS cells have a higher spine density comprising of more thin spines than IB cells and conversely, IB cells have a lower spine density but a greater proportion of mushroom spines.

**Table 1.**
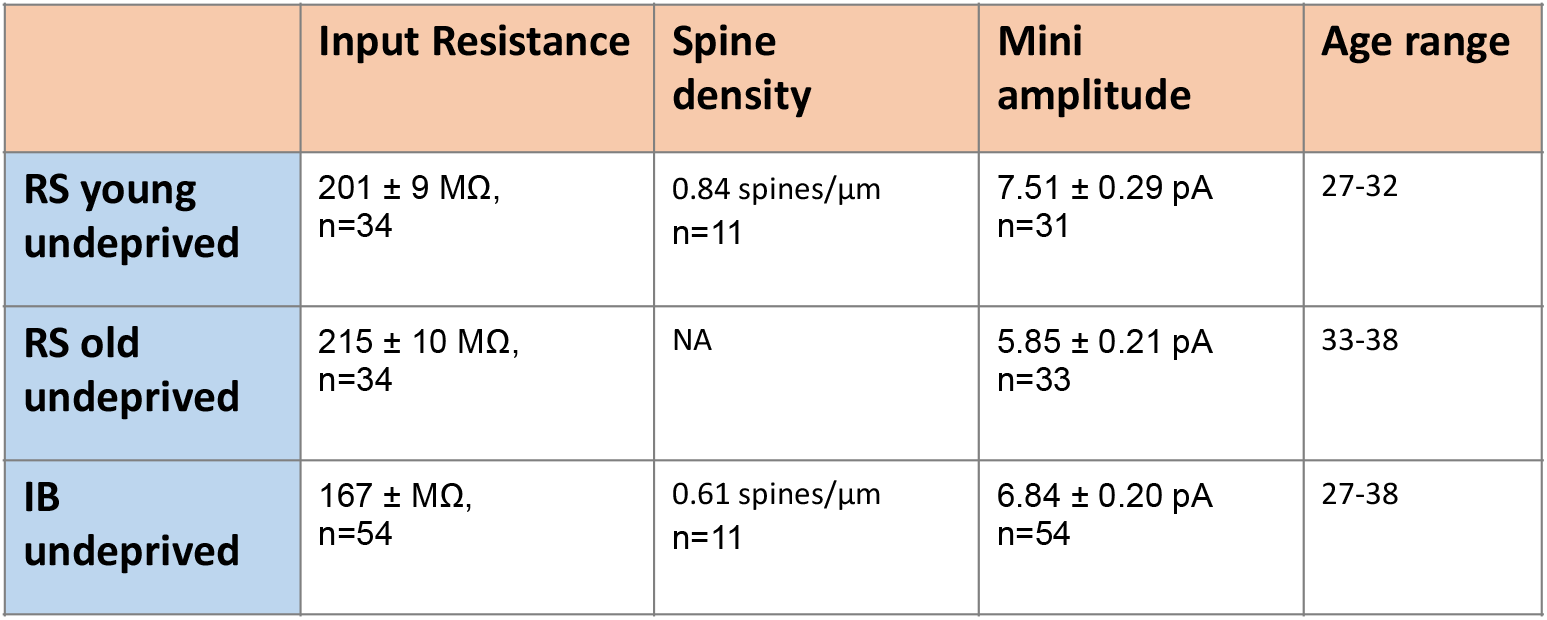
Average statistics for control conditions. Input resistance, spine density and mEPSC amplitudes are shown for the two RS cells age groups (young=27-32 days; old =33-38 days) and the IB cells (P26-38).

### Development

During the critical period for ocular dominance plasticity (P19-32) (23), mEPSC amplitudes in L5 RS cells decreased with age, whereas those for IB cells remained constant over time (Fig. 2; Table 1). We compared mEPSCs in two age groups, P27-32 and P33-38. A 2-way ANOVA showed a strong effect of age on mEPSC amplitude (F_(1,1)_=2.81, p<0.006) and an interaction between age and cell type (F_(1,1)_ =12.43, p<0.006). Post hoc t-tests showed that this was because earlier in the critical period the average mEPSC amplitude for RS cells was higher (at 7.5pA) compared with at the end (5.8pA) and the two values were highly significantly different (t_(62)_=4.97, p<0.001), whereas mEPSCs for IB cells were similar at the two ages (6.7pA and 6.9pA) and not statistically different (t_(53)_=0.21, p=0.65). In corroboration, we also found the linear regression fit for a plot of mEPSC amplitude versus age had a negative slope significantly different from zero for RS cells (F_(1,9)_=13.2, p<0.01) and correlated with the data (R^2^ = 0.59), whereas the regression fit for IB cells was flat (F_(1,9)_=0.71, p<0.42) and uncorrelated with the data (R^2^ = 0.07).

**Figure. 2.**
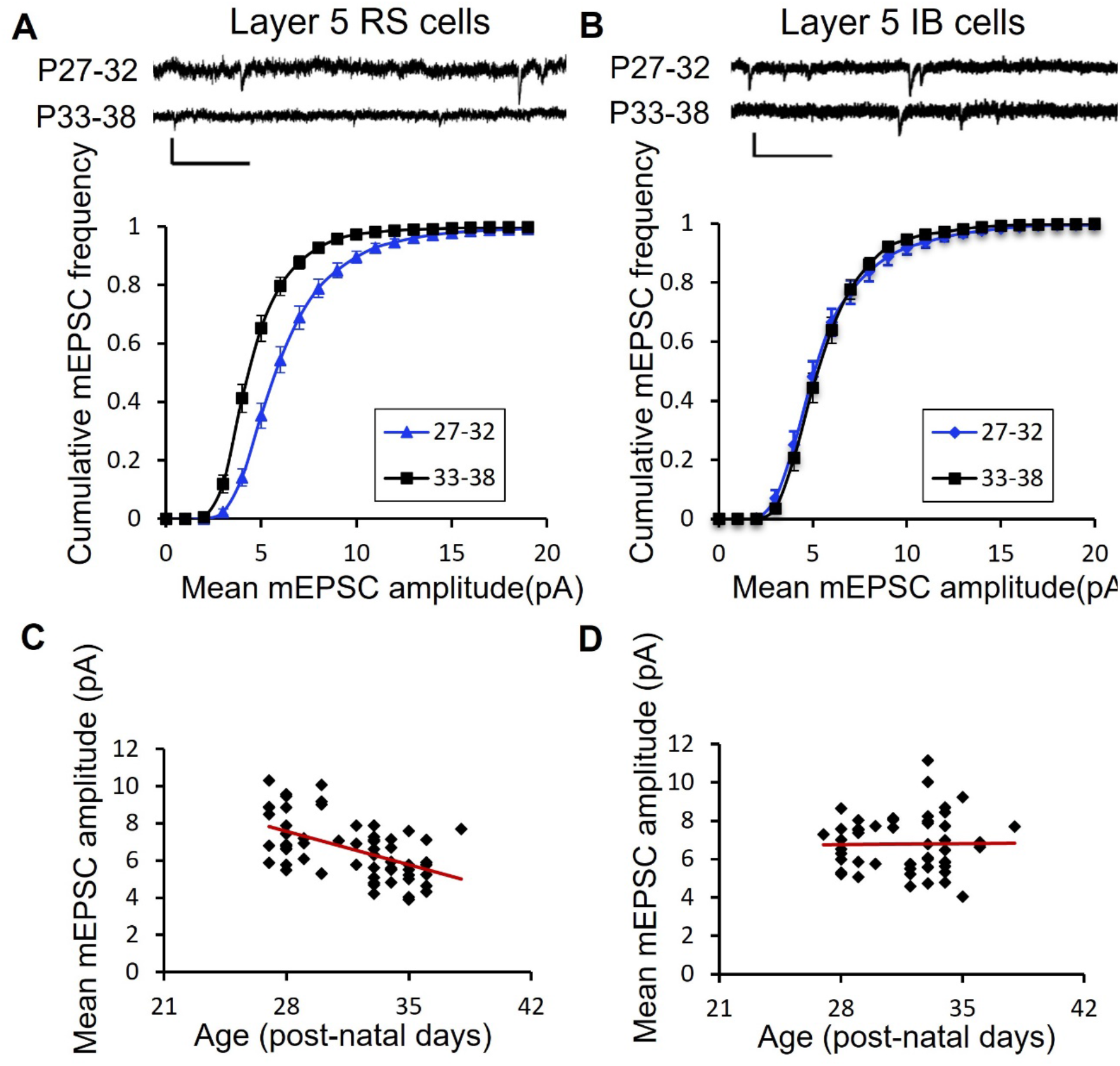
Synaptic development of layer 5 RS and IB neurons. **A:** Example mEPSC traces and cumulative distribution functions for RS cells in the P27-32 (blue) and P33-38 (black) age group (scale bars 10pA and 250ms). The two age groups are significantly different (see Results). **B:** Example mEPSC traces and cumulative distribution functions for IB cells in the P27-32 (blue) and P33-38 (black) age group (scale bars 10pA and 250ms). The two age groups are not significantly different (see Results). **C:** RS cells show a decrease in synaptic efficacy between P27 and P38 whereas **D:** IB cells show no change over this period (see text for statistical significance).

The changes in mEPSC amplitude with age in RS neurons could not be attributed to changes in input resistance (P27-32: R_in_ = 245 ± 16.8MΩ; P33-38: R_in_ = 251± 18.5MΩ; t_(51)_=0.87, p=0.38; Table 1). We tested whether the frequency of the mEPSCs changed with age and found they did not, either for IB (K-S test, D=0.08, n=101 p>0.1) or RS cells (K-S test, D=0.03, n=101 p>0.1). We conclude that L5 RS cells continue to develop throughout the critical period, reducing their synaptic amplitude, while IB cells have either already completed this stage of development or do not develop this way at all. Therefore, in the results described below, for RS cells we have compared the experimental data against closely age-matched undeprived controls, while the IB cells are compared across the P27 to P38 time period.

### Monocular Deprivation

Monocular deprivation (MD) is the classic method for investigating plasticity in the visual cortex (11, 23, 24) and has been shown to induce Hebbian and homeostatic components of plasticity (25). However, studies have mainly been directed at layer 2/3 and layer 4 cells and, where studies have addressed plasticity in layer 5, they have usually not differentiated between RS and IB neurons [but see (26)].

For RS neurons located contralateral to the closed eye and in the binocular zone (see Methods), we found that MD caused a rapid depression in mEPSC amplitude after 12 hours that was sustained to 3 days, but which showed a homeostatic rebound to baseline values at 5 days (Fig. 3). An ANOVA showed a strong effect of deprivation on age-matched mEPSC amplitude (F_(4,4)_ = 6.45, p<0.0001) and post-hoc t-tests showed that only the 12 hour and 3 day time points were different from undeprived age-matched controls (12h, t_(34)_=2.87, p<0.01; 3d, t_(38)_=3.2, p<0.005).

**Figure. 3.**
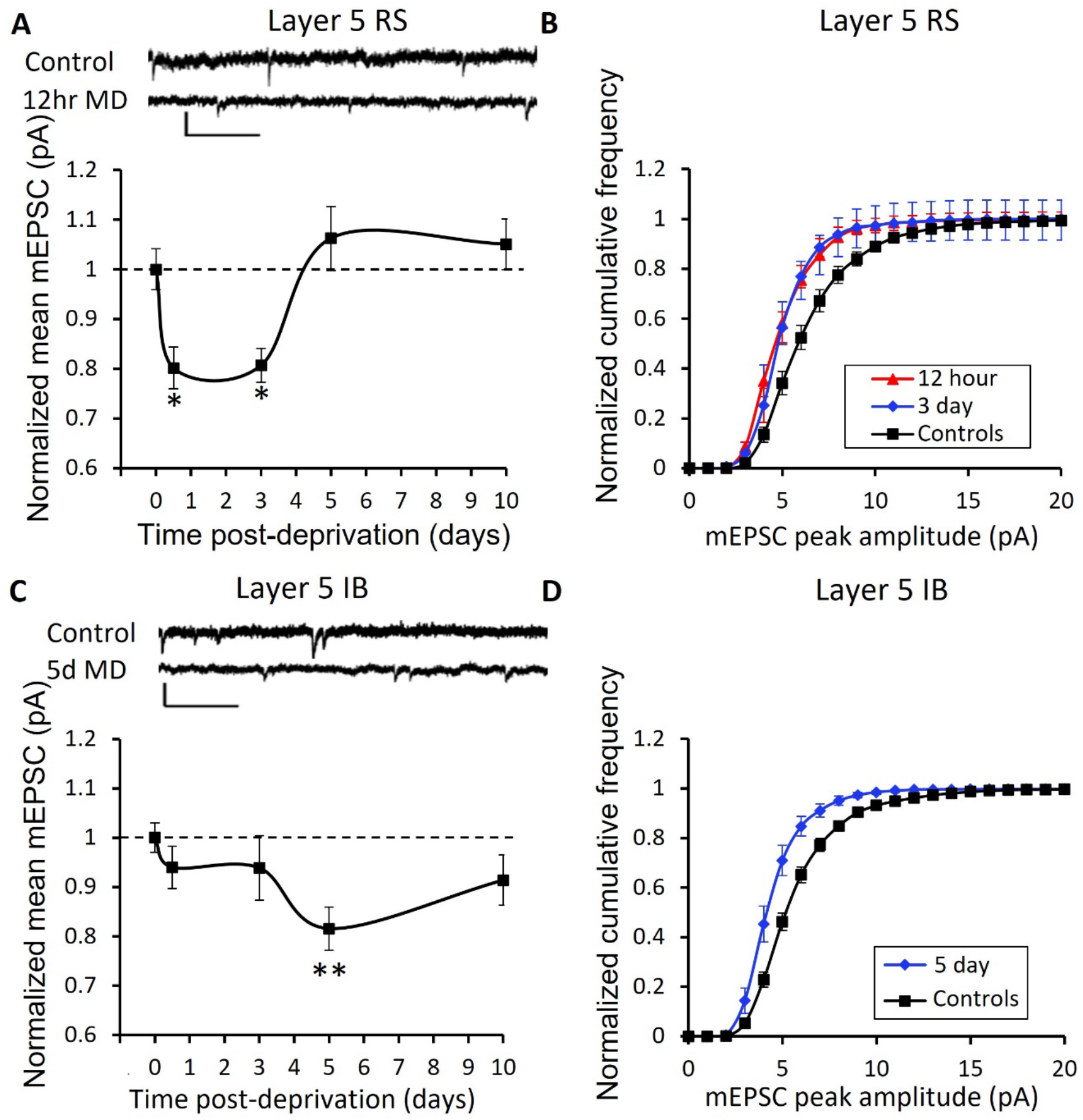
Effect of monocular deprivation (MD) on synaptic strength in Layer 5 RS and IB cells. **A:** Example mEPSCs and time course of change in RS cells’ mEPSCs following monocular deprivation (scale bars 10pA and 250ms). **B:** Cumulative distribution functions (CDFs) for control and the two significantly depressed time points shown in ‘**A’** at 12 hours (red) and 3 days (blue). **C:** Example mEPSC traces and time course of changes in IB cells’ mEPSCs following monocular deprivation (scale bars 10pA and 250ms). **D:** Cumulative distribution function for the 5-day time point (blue) compared to control (black). *p<0.05, **p<0.01. For CDFs of all timepoints see Extended Data Fig. 3.

The IB neurons showed a delayed effect of deprivation in comparison with the RS neurons. The mEPSC amplitudes at the 5 day time point showed depression compared to controls (Fig. 3). An ANOVA showed an effect of deprivation (F_(4,96)_ = 2.5, p<0.05), but none of the deprivation time points were distinguishable from controls except the 5 day time point (t_(65)_=2.87, p<0.01). A homeostatic rebound occurred between 5 and 10 days (Fig. 3). The IB cells therefore behaved similarly to the RS cells, but delayed in time both in depression onset and rebound.

We also tested to see whether the frequency of mEPSCs was affected by MD (Extended Data Fig. 4) and found that it had no effect on IB cells at any time point (Kruskall Wallis (KW) test on average inter-event interval per cell, χ^2^ = 4.46, df=4, p=0.35), but did affect RS cells (KW test on average inter-event interval per cell, χ^2^ = 9.98, df=4, p<0.05). Post hoc tests revealed that inter-event intervals increased at both the 12 hour and 5 day time points (Wilcoxon test, α=0.05). The reduction in mEPSC frequency at 12 hours would be expected to exacerbate the effect of the reduced mEPSC amplitude. However, the reduced frequency at 5-day timepoint would tend to offset the effect of the mEPSC’s return to baseline values at this time point (Fig. 4A).

**Figure. 4.**
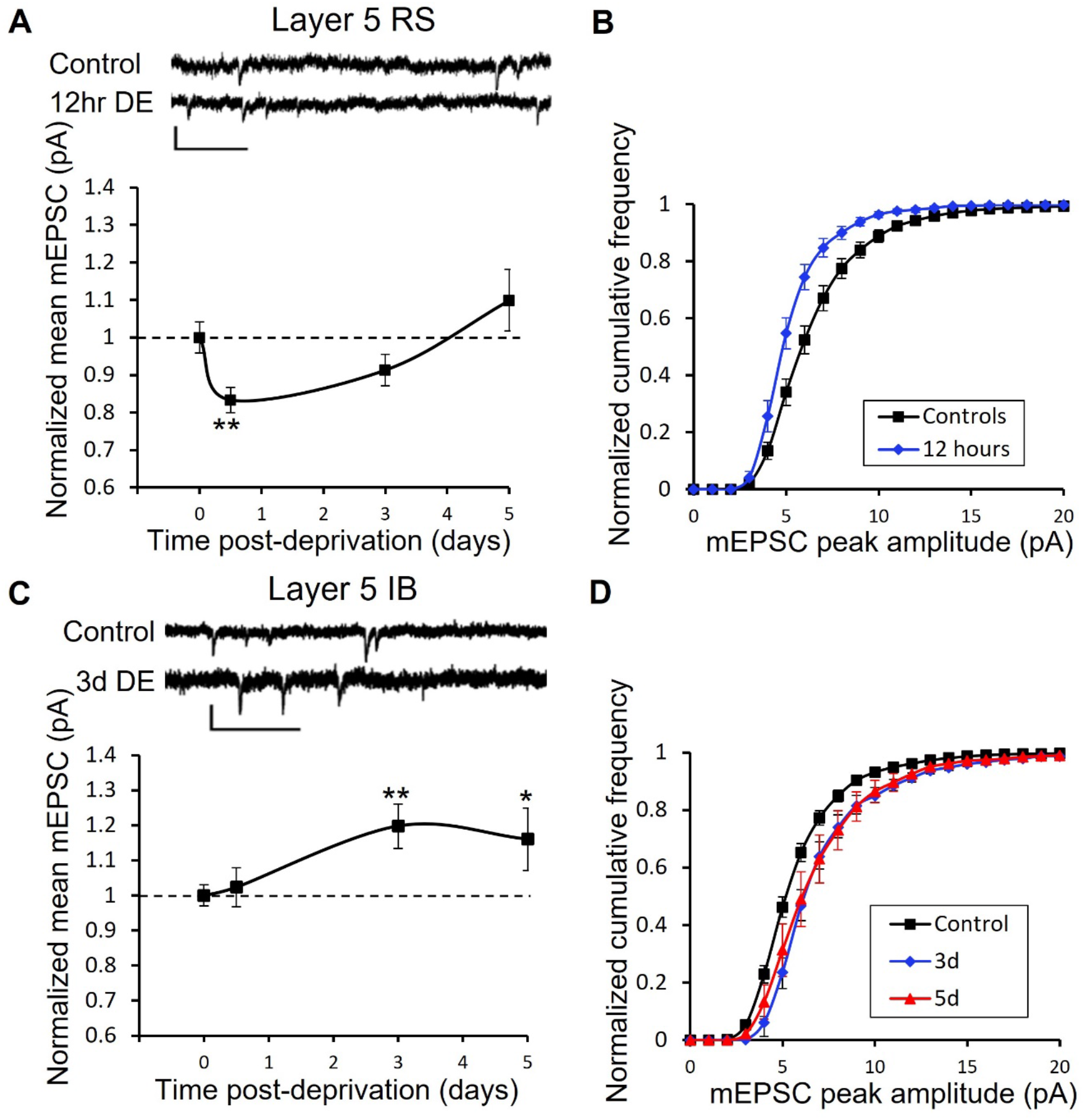
Effect of dark exposure (DE) on synaptic strength in Layer 5 RS and IB cells. **A:** Example mEPSC traces and time course of change in RS cells’ mEPSCs following dark exposure (scale bars 10pA and 250ms). **B:** Cumulative distribution functions for control (black) and the 12 hour (blue) significantly depressed time point shown in ‘**A’** (scale bars 10pA and 250ms). **C:** Example mEPSC traces and time course of changes in IB cells’ mEPSCs following dark exposure. **D:** Cumulative distribution function for the 3 day (blue) and 5 day (red) time points. *p<0.05, **p<0.01. For all CDFs see Extended Data Fig. 5.

In addition, we tested whether the input resistance (R_in_) changed during the deprivation. We found that 12 hours of MD increased the input resistance in both RS and IB cells, but it returned to control values for all other time-points. A 2-way ANOVA showed an effect of cell-type (F_(1,1)_=16.03, p<0.001) and deprivation (F_(4,4)_=2.76, p<0.05) but no interaction. IB cells had lower input resistance than RS cells (IB cells: 167+7.5MΩ versus RS cells: 208+6.7MΩ) (Table 1), and the difference was highly significant (t_(210)_=4.0, p<0.001). Post-hoc t-tests also showed that R_in_ increased only at the 12 hour time-point (t_(210)_=3.3, p,0.002) independent of cell type. The increase in R_in_ at 12 hours may have some effect in offsetting the synaptic depression at 12 hours in RS cells, but because it is not sustained beyond this time-point it would not affect the visual responses at 3 days of MD.

### Dark Exposure

Dark exposure (DE) has been used as a model for inducing synaptic scaling in layer 2/3 visual cortical neurons (16). In this study, we wanted to determine whether Layer 5 RS and IB cells both underwent synaptic scaling and to what extent the process was homeostatic.

A 2-way ANOVA showed an effect of cell type (F_(1,1)_=11.85, p<0.001), duration of DE (F_(3,3)_=4.05, p<0.01) and an interaction between the two (F_(3,3)_=3.95, p<0.02). Analysing the effect of DE on RS cells separately, post hoc t-tests with age matched controls showed that DE caused rapid depression in mEPSC amplitudes after 12 hours (t_(46)_=2.76, p<0.01), that returned to baseline after 3 days (t_(45)_=1.05, p=0.29), similar to the effect of MD on RS cells but with a faster homeostatic return to baseline (Fig. 4).

The effect of dark exposure on IB neurons was fundamentally different from its effect on RS cells in that it caused clear potentiation, rather than depression and homeostatic rebound (Fig. 4). Post-hoc t-tests showed potentiation was significant at the 3 day (t_(65)_=2.85, p<0.01), and 5 day time points (t_(63)_=2.44, p<0.02).

We also tested whether dark exposure altered the inter-event intervals of mEPSCs (Extended Data Fig. 6) and found that it did not in IB cells at any time point (KW test on average inter-event intervals, χ^2^ = 3.4, df=3, p=0.32) but did have an effect on RS cells (KW test, χ^2^ = 11.9, df=3, p<0.01). Post-hoc tests showed that inter-event intervals were specifically affected at the 5 day time point (Wilcoxon test, Z=3.2, p<0.002). The reduction in mEPSC frequency might therefore offset the recovery of the mEPSC amplitude to baseline values at 5 days (Fig. 5A) to some extent.

**Figure. 5:**
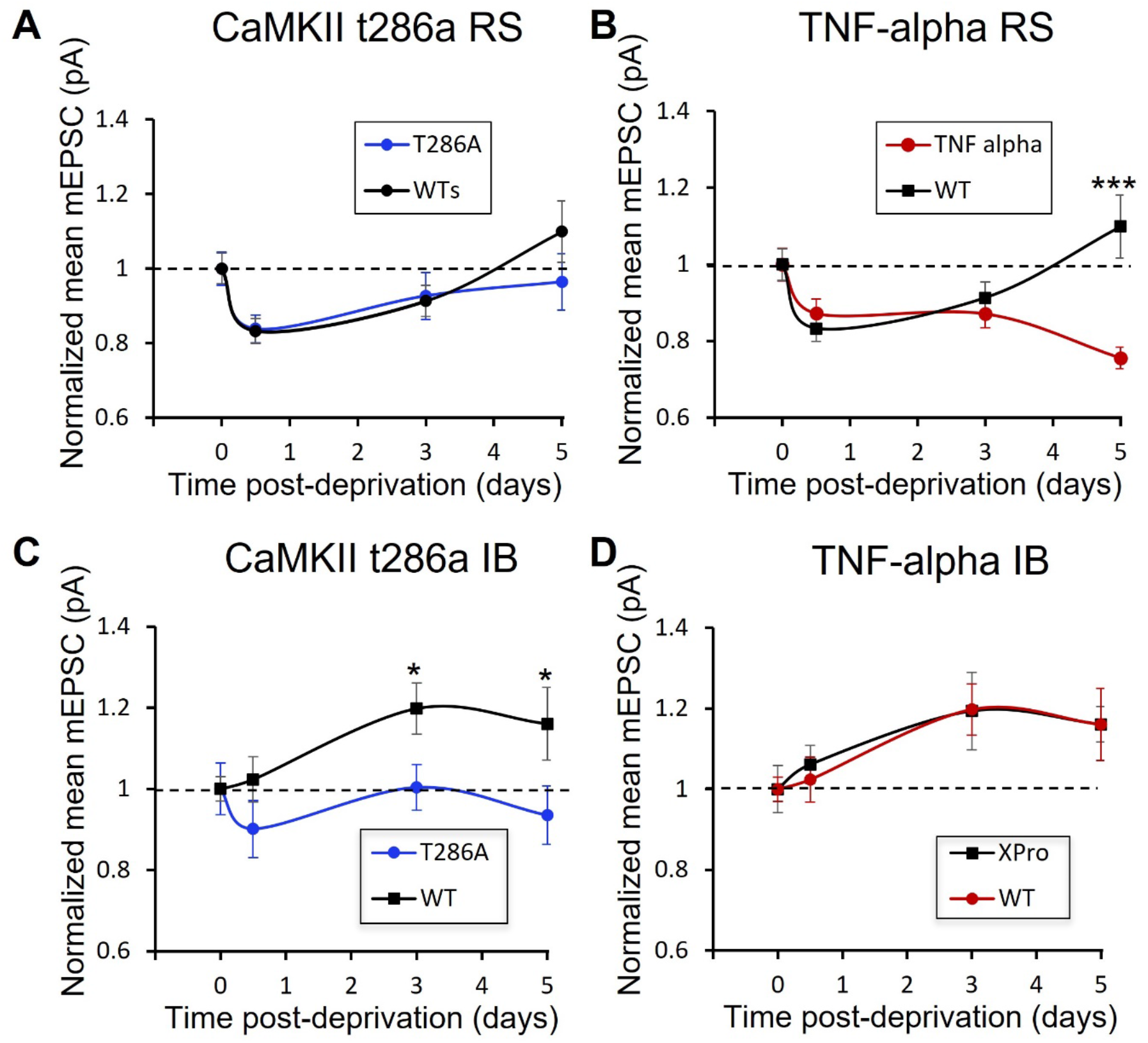
RS and IB neurons show different plasticity mechanisms during dark exposure. **A:** Depression and homeostatic rebound to baseline values in wildtypes (black) and CaMKII t286a mutants (blue) are similar in RS cells whereas **B:** homeostatic rebound does not occur in wild-type RS cells treated with TNFα inhibitor (red). **C:** Potentiation does not occur in IB cells of CaMKII-t286a mutants (blue), however **D:** potentiation is unaffected by treatment with TNFα inhibitor (red).

### TNFα and αCaMKII-autophosphorylation dependence of plasticity

The different responses of RS and IB cells to dark exposure suggest that different plasticity mechanisms may underlie the effects. The RS cells show depression followed by homeostatic rebound to control values, which is a signature of homeostatic plasticity, previously found to depend on TNFα (9, 16). We tested to see whether plasticity was TNFα dependent using the injectable TNFα dominant negative peptide XPro1595, which combines with endogenous extracellular TNFα and renders the resultant trimeric agonist ineffective at binding at TNFα receptors (27).

We found that XPro1595 had no effect on the depression induced in RS cells by dark exposure, but that it prevented the homeostatic rebound in the mEPSC amplitude that occurred between 3 and 5 days in untreated animals (Fig. 5). A 2-way ANOVA showed an effect of dark exposure and XPro1595 on the mEPSC amplitude and an interaction between the two (F_(3,3)_=5.7, p<0.005). The interaction arose from the 5 day timepoints being different between XPro1595 treated and untreated cases (t(24)=, p<0.001) while none of the other timepoints were different (t-test, α=0.05). In contrast to the effect on RS cells, XPro1595 had no effect on IB cells (Fig. 5D). These results suggest that the rebound observed in RS cells toward baseline response levels is TNFα-dependent while the potentiation seen in IB cells away from the baseline values is not.

Previous studies in barrel cortex had shown potentiation in layer 5 IB cells to be dependent on αCaMKII-autophosphorylation (9). CaMKII-t286a mutants have calcium sensitive CaMKII, but lack the ability to autophosphorylate at the threonine 286 site. The autophosphorylation step is necessary for LTP to be sustained *in vitro* (28–30) and is also important for dendritic spine enlargement *in vivo* (31). We therefore tested whether the potentiation of the mEPSCs produced in IB cells was CaMKII-autophosphorylation dependent.

We found that potentiation produced by 3-5 days dark exposure was absent in layer 5 IB cells in mice carrying the mutation preventing CaMKII-autophosphorylation (CaMKII-t286a mice) as shown in Fig. 5C. A two way ANOVA showed an effect of genotype (F_(3,3)_=4.7, p<0.005) and post hoc t-tests showed that this was because the 3 and 5 day timepoints were different in wild-type mice compared with the CaMKII-t286a point mutants (t_(22)_=, p<0.05). In contrast the depression and homeostatic response of the RS cells recorded in these mutants was the same as found in wild-type animals. A 2-way ANOVA showed no difference between RS cells response to dark exposure in wild-types and mutants, only an effect of deprivation (F_(3,3)_=5.09, p<0.005). These findings therefore show that the potentiation seen in IB cells depends on the same molecular mechanism as LTP, experience-dependent potentiation and experience-dependent dendritic spine enlargement, whereas the depression and rebound homeostatic potentiation seen in the RS neurons does not.

### Morphological correlates of changes in mEPSCs

We looked at the morphology of dendritic spines to see if we could identify structural changes that corresponded to the electrophysiological changes we observed with dark exposure. Consistent with other studies, we found that spine head sizes were log-normally distributed. In IB cells, we found that there was an effect of dark exposure on spine head size and that in the undeprived case, spine head sizes in apical and basal dendrites were significantly different. A 2-way ANOVA showed an effect of dendritic location (F_(1,1)_=12.6, p<0.001) and deprivation on spine head size (F_(2,2)_ =5.7, p<0.01). Post-hoc test showed that spine head sizes were far smaller on basal dendrites when compared with apical dendrites (t_(21)_=12.0, p<0.003). Spine head size increased after just 12 hours of deprivation on basal dendrites (compared with control underived cases, t_(20)_=15.8, p<0.0001) before regressing back toward control values after 3 days (Fig. 6A). What appear to be changes in apical dendritic spine size with deprivation were not statistically significant (t_(20)_=2.4, p=0.13). An ANOVA did not reveal any differences in spine head size in RS neurons; although it appeared that apical dendrites may have enlarged at 12 hours (Fig. 6B), this was not statistically significant (t_(18)_=2.1, p=0.16).

**Figure. 6.**
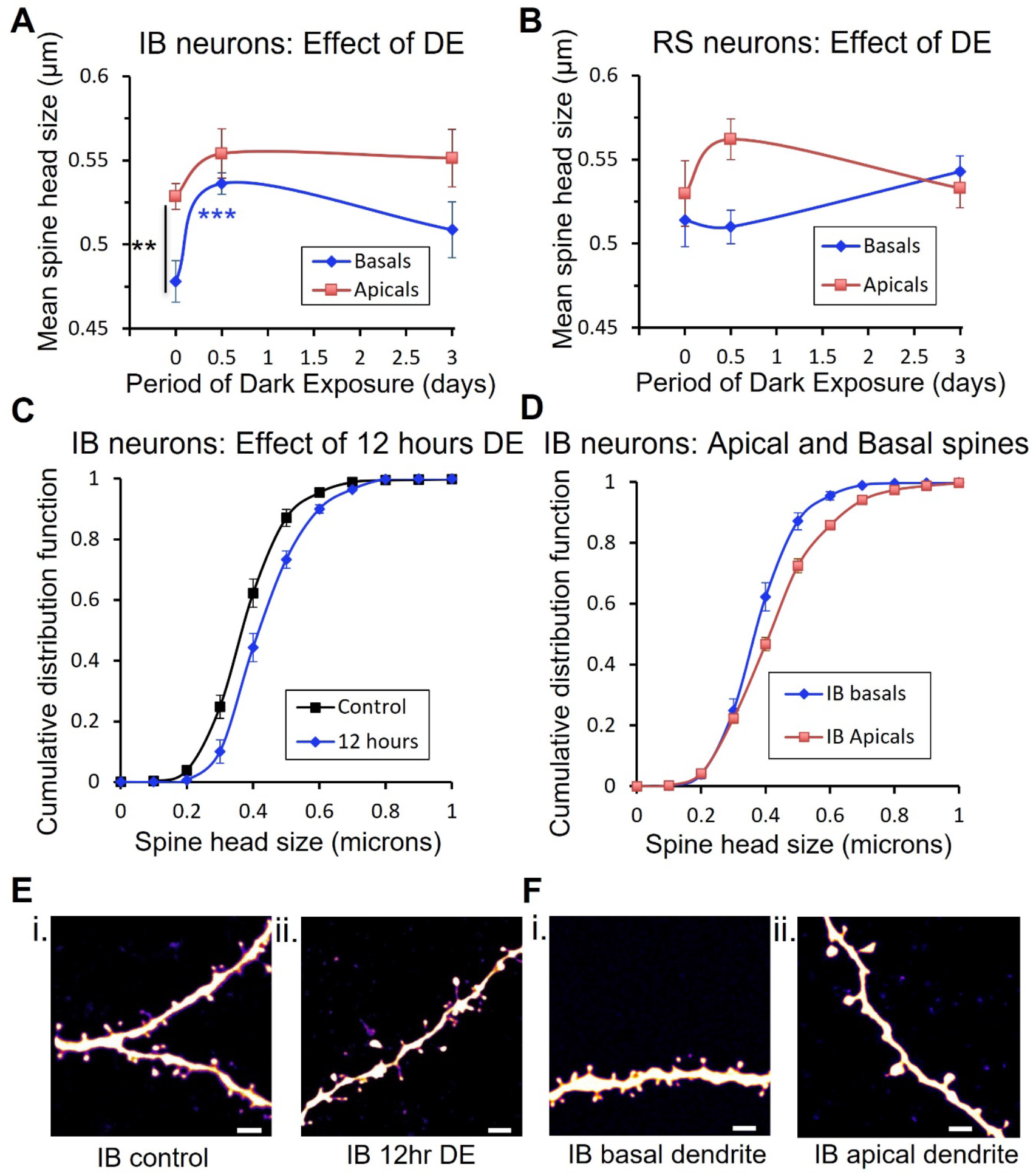
Changes in spine head diameter following dark exposure. **A:** In IB cells, spine head size increases for spines on basal dendrites after 12 hours of dark exposure. **B:** No significant changes in spine head size could be detected in RS cells. **C:** Cumulative distribution functions for basal dendritic spines from IB cells show the increase in IB spine head size after 12 hours DE (blue) is uniform across the population of spines. **D:** Spine head sizes show a scaling difference between apical (red) and basal (blue) spines from IB cells. **E (i):** Example dendritic spines on basal dendrites of IB neurons from a control animal **E(ii):** basal dendrites of IB neurons from an animal after 12hr DE. **F(i):** basal dendrites of an IB neuron from a control animal. **F(ii):** apical dendrite of an IB neuron from a control animal. Scale bar in spines images 2μm.

We found that apical dendritic spine sizes were only different from basal dendritic spines in undeprived cases (Fig. 6A) and that dark exposure negated the difference. The increase in spine head size after 12 hours appeared to be due to a small (60nm) and relatively uniform rightward shift in spine size values Fig. 6C. The difference in spine sizes between apical and basal dendrites (Fig. 6D) is likely to be related to distance dependent scaling (32), which is thought to compensate for dendritic EPSP attenuation distant from the soma.

The changes in spine head size occurred in the absence of any overall change in spine density on the basal dendrites of RS or IB cells (Extended Data Fig. 7). However, we did observe an increase in the spine density on apical dendrites of IB cells (but not RS cells) after 12 hours of deprivation (2-way ANOVA; interaction between spine location and deprivation, F_(2,2)_=4.3, p<0.02).

Our structural plasticity findings therefore show that, for IB cells, changes in basal spine head size and apical spine density occur after 12 hours of deprivation, before any electrophysiological changes are detectable in the mEPSC amplitudes (compare Fig. 4C and 6A). Conversely, when the electrophysiological changes are seen in IB cells at 3 days, the structural changes have decreased slightly. For RS cells, we did not find any overt structural changes in spine density or spine head size whatsoever that might correlate with decrease in mEPSC amplitudes seen after 12 hours deprivation.

## Discussion

### Generality of RS/IB dichotomy

One of our objectives was to test whether the distinction between IB and RS synaptic plasticity properties can be generalised across cortical areas. Despite the very different nature of visual and somatosensory information, we found that IB and RS cells exhibited the same differences in plasticity mechanisms in visual cortex as they did in somatosensory cortex. In both cortical areas, IB cells showed potentiation beyond baseline values; in visual cortex in response to dark exposure (Fig. 4C) and in barrel cortex in response to chessboard pattern whisker deprivation (9). The IB cell potentiation was also dependent on CaMKII-autophosphorylation in both areas and not dependent on TNFα (in either cortical area). Similarly, RS cells in both cortical areas showed synaptic depression in response to sensory deprivation, independent of the type of deprivation; monocular deprivation or dark exposure for visual cortex (Fig. 3A, 4A), and chessboard pattern deprivation or all-whisker deprivation in the barrel cortex (9). In both cases, the homeostatic rebound in the response of the RS cells followed a similar time-course in the two cortical areas (of approximately 3 days) and was dependent on TNFα and not CaMKII-autophosphorylation.

### Interpreting visual cortex plasticity experiments

Our studies show that the time course and direction of plasticity are different in RS and IB cells, which needs to be considered when understanding the effect of monocular deprivation and/or dark exposure. In fact, a different interpretation emerges if we ignore the distinction between cell types. Extended Data Fig. 8 shows a plot of the time course of synaptic strength during dark exposure and monocular deprivation disregarding celltype. Under these circumstances, the mean EPSC amplitudes appear to show a small depression and return to baseline following either form of deprivation; for dark-exposure the RS and IB cell EPSC amplitudes move in opposite directions cancelling one another out so that the only time point that differs from the aggregated control data is the 5 day time point that shows a small potentiation of 12% (t_(128)_=5.8, p<0.02). For monocular deprivation, the combined RS and IB populations show depression (maximum at 3 days, 14% reduction, t_(124)_=8.6, p<0.005) as with the individual cell types, but again the effect of the RS and IB cells moving in opposite directions between 3 and 5 days cancel one another out such that the homeostatic rebound is only complete after 10 days whereas for RS cells this really occurs in 5 days.

The similarity of the RS cells’ reaction to monocular deprivation and dark exposure can probably be explained most easily by their strong contralateral bias (33), since we recorded neurons in the hemisphere contralateral to the monocular deprivation. The RS cells were therefore deprived of their dominant contralateral input during MD. While the ipsilateral input was additionally deprived during dark exposure, which would not affect RS cells much more, as they are strongly contralateral biased.

### Cellular diversity in plasticity mechanisms

While the RS cells show a classic depression followed by TNFα dependent synaptic scaling, IB cells show a mechanism more closely related to LTP, judging by its dependence on CaMKII-autophosphorylation. Recent studies on the effects of dark exposure in layer 2/3 cells in visual cortex have shown that plasticity occurs via NMDA receptors rather than synaptic scaling (34). This is consistent with experiments showing NR2B up regulation following dark rearing (35). Similarly, blocking NMDA receptor function by acute knock-out prevents the increase in mEPSC levels seen in visual cortex L2/3 cells (36). This type of L2/3 NMDA dependent mechanism seems to be similar to the one we describe for L5 IB cells, since NMDA receptors induce plasticity via calcium and CaMKII-autophosphorylation (to produce LTP), promote new spine survival and enlargement of pre-exiting spines (28, 29, 31).

However, the same mechanism is not triggered in layers 5 RS cells during dark exposure; they do not show potentiation and their homeostatic recovery is not CaMKII dependent. The same can be said for RS cells in barrel cortex (9). The situation appears to be more complicated for layer 2/3 cells, which are mostly RS (37) and show both Hebbian and synaptic scaling properties in visual (16) and somatosensory cortex (38).

### Relationship between structural and functional changes

In this study, we discovered a number of previously unreported differences between IB and RS cells. While IB cells showed mature synaptic properties at the start of the critical period, RS cells in layer 5 matured during the critical period. Furthermore, the two cell types showed different spine morphologies, with IB cells having a greater proportion of mushroom spines and RS cells having a higher proportion of long thin spines than IB cells.

In most cases we did not find clear correlations between structural and functional changes, however, in the case of DE we found that the structural changes in IB cells preceded the functional changes; spine enlarged after just 12 hours dark exposure whereas the mEPSC amplitude was on average, unchanged at this time point. The functional increase in mEPSC amplitude was observed at 3 days, by which time the spine size had fallen back from its peak (at 12 hours). These observations are consistent with the findings of two separate studies, each of which reported one of the effects. First, Barnes et al., (2017) detected an increase in spine head size just 8 hours after eye-enucleation (17). Second, in a separate study, recordings from layer 5 in animals receiving retinal lesions showed no increase in mEPSC amplitude at 6 and 18 hours; instead, increases only became apparent after 24 hours (39). Taken together with the present study, it would appear that an increase in spine head size precedes a functional increase in mEPSC amplitude.

Studies in hippocampal cultured cells have also demonstrated that the first change following NMDA receptor activation is an increase in spine size, which precedes an increase in AMPA receptor surface expression (40). Both changes in structure and function are controlled by CaMKII as evidenced by *in vitro* (41) and *in vivo* studies (31, 42). Crucially, we find that DE does not produce functional changes in mice lacking CaMKII-autophosphorylation, which indicates that the spine enlargement mechanisms described in these methodologically different studies (39–41) are related to our observations in IB cells.

## Materials and Methods

### Subjects and sensory deprivation

All procedures were approved under the Animal (Scientific Procedures) Act 1986. A total 268 animals were used in the study across different age groups, genotypes, deprivation time points, cell types, visual deprivation group (monocular deprivation or dark exposure) and treatment group (XPro 1595) (see Supplementary Tables 1 and 2). All the mice used were either WT C57Bl/6J (Charles River) or homozygous T286A homozygous mutants on the C57Bl/6J background. The control mice were housed in standard housing conditions of 12 hrs light/dark cycle. All the animals used were 26-38 days old. Visual deprivation was started at the age of 26-31 days. For dark exposure experiments, two mice were housed together in the dark room at a time. For monocular deprivation experiments the eyelid of one eye was sutured under isoflurane anaesthesia (6-O non-absorbable sutures, Mersilk-Ethicon, J&J) and topical antibiotics applied (1.0% Chloramphenicol 1.0% w/w Eye ointment, Martindale Pharma). The sutures were checked every day to monitor closure and animals were not used if gaps developed.

A subset of animals was injected with XPro-1595, 1mg per Kg of animal’s body weight. XPro-1595 was injected at least 12hrs before the electrophysiological recordings, or start of dark exposure. In case of 5d DE XPro-1595 was injected twice - 12hr before the start of DE and on 3rd day of DE. Control animals corresponding to XPro-1595 injected DE animals were also similarly injected with XPro-1595.

### Surgeries for injection

For labelling neurons projecting to various targets retrograde latex beads (RetroBeads, Lumafluor inc., NC, USA) were injected in the putative target sites. For labelling contralateral visual cortex projecting neurons, retrobeads (100 nl of 4X diluted solution at 10nl/min) were injected at two locations in visual cortex layer 5 (−3.8 mm AP from Bregma, 2.5 mm ML from midline, 0.7 mm DV from Pia and −3.5 mm AP from Bregma, 2.3 mm ML from midline, 0.7 mm DV from Pia). To label neurons projecting to the Superior colliculus, retrobeads were injected at (−3.9 mm AP from Bregma, 0.1 mm ML from midline, 2.2 mm DV from Pia) with the injection needle tilted at 25° from the midline. The injection needle was held at the injection location for 10 minutes after injection to make sure there was no backflow of the beads in superficial layers.

### In vitro electrophysiology

Mice were killed by decapitation. The brain was quickly removed and immediately placed in ice cold slicing solution (in mM: 108 Choline-Cl, 3 KCl, 26 NaHCO3, 1.25 NaH_2_PO4, 25 D-glucose, 3 NaPyruvate, 1 CaCl_2_, 6 MgSO_4_, 285 mOsm, bubbled with 95% O_2_ 5% CO_2_). 350μm thick coronal slices were cut in ice cold slicing solution using a vibrating microtome (Microm HM650V). Slices were then transferred to a holding chamber containing normal ACSF (in mM: 119 NaCl, 3.5 KCl, 26 NaHCO_3_, 1 NaH_2_PO_4_, 2 CaCl2, 1 MgSO_4_, 10 D-glucose, 300 mOsm bubbled with 95% O_2_ 5% CO_2_). Slices were incubated at 37°C for 30 minutes and then returned to room temperature before recording. The area of binocular visual cortex was identified using mouse brain atlas by Paxinos (43). We recorded mEPSCs from the hemisphere contralateral to deprived eye within the binocular zone. Layers 5A and 5B were identified under bright field illumination using differential interference contrast on an Olympus BX50WI microscope, guided by the neuronal density and morphology of the cells. Whole cell voltage/current clamp recordings were performed using borosilicate glass electrodes (4-7 MΩ) filled with a potassium-gluconate based solution (in mM: 110 K-gluconate, 10 KCl, 2 MgCl_2_, 2 Na_2_ATP, 0.03 Na_2_GTP, 10 HEPES, pH 7.3, 270 mOsm). In a subset of experiments biocytin (Sigma, UK) was added to the electrode filling solution at a concentration of 5mg /ml to enable cells to be morphologically characterised.

Pyramidal neurons were classified into IB and RS neurons based on their spiking properties at threshold. Briefly, the neurons were classified as IB neurons, if the first action potentials fired by the neuron at threshold were in the form of one or multiple bursts, otherwise the neurons were classified as RS neurons. A burst was defined as at least 2 action potentials firing at a frequency higher than 100 Hz as described earlier (44). After the characterisation of active properties of the neurons, the slices were perfused with ACSF containing a cocktail of drugs consisting of 50 *μ*M DL-AP5, 10*μ*M picrotoxin and 1*μ*M TTX. The mEPSCs were recorded in the voltage clamp configuration. Recordings were made using an Axon Multiclamp 700B, digitized with a CED Micro 1401 controlled with CED Signal software. Miniature EPSCs were analysed with Axograph software using a template-matching method (see data analysis below).

### Dendritic spine imaging

After electrophysiological recordings, slices containing biocytin filled neurons were fixed overnight at 4°C in 100 mM phosphate buffered saline (PBS, pH 7.3) containing 4% paraformaldehyde (BDH, USA). Thereafter, slices were washed with PBS and incubated in PBS supplemented with 1% Triton X-100 and 0.2% streptavidin Alexa Fluor 488 conjugate (Invitrogen) at 4°C for 18hrs. After further washing with PBS, cells were imaged under a 2-photon microscope (Prarie Systems) using a mode-locked Ti:sapphire laser (Chameleon Vision S, Coherent) to generate two-photon excitation (900nm), and emission wavelengths were band-passed between 525-570 nm (including an IR filter in the light path). A 10x objective lens was used to image gross cell morphology while a 60x objective lens was used for detailed dendritic and spine morphology. For spine morphology 60X objective was used and images were digitally magnified at 4.02 times. The complete dendritic morphology was measured for each filled neurone, while spine morphology was recorded for a subset of the dendrites, which fully described the neurone’s dendritic arborisations (basal dendrites, apical obliques and apical tuft). All the imaged spines were located on secondary or tertiary dendritic branches. Image stacks were collapsed into z-stacks (Image J) which in turn were stitched together (Photoshop, Adobe) to produce a detailed 2D dendritic profile of the neuron (Figure 1A). The distance of the soma from the pia, length of apical dendrites, sites of bifurcation and the horizontal spread of the dendrites were all measured using Image J (NIH, USA).

### Morphological analysis

The neurons’ dendritic fields were analysed using Sholl analysis. We counted the number of occasions that the dendrites crossed the Sholl shells (radial interval = 30μm) at increasing distance from the soma. The Sholl shells were centred at the soma for basal and apical oblique dendrites, while for apical tuft they were centred at the main bifurcation of the apical dendrite. We also measured the distance of the cell bodies from the pial surface, the length of the apical dendrites and the distance of apical bifurcation from the soma.

### Measurements of spine morphology

Dendritic spines were analysed as described earlier (31). In brief, Image stacks were first deconvolved using Fiji Deconvolution Lab plugin and the spines were measured using ImageJ (NIH, USA). Only structures that were protruding at least 0.4 μm from the dendritic branches were counted as spines (Holtmaat et al., 2009). Spines were classified based on head size, neck width, and neck length measurements as described earlier (Holtmaat et al., 2009). Briefly, spines with head size: neck width ratio >1.15 and a neck length < 0.9 μm were classified as Mushroom spines, while thin spines had head size: neck width ratio >1.15 and a neck length >0.9 μm. Stubby spines had a neck length < 0.9 μm and a head size: neck width ratio <1.15. We also saw a smaller number of filopodia, which have neck length >0.9 μm while head size: neck width ratio <1.15. Filopodia were not included in the spine analysis unless stated, as with the spine classification sections. At least 60 spines were measured from each type of dendrite (apical, oblique or basal) for each neuron. Since the percentage of filopodia in the total spine population was very low, filopodia were excluded from the final analysis except for the quantification of spine types (as detailed in the results section).

### Data analysis and statistics

The mEPSCs were analysed using Axograph X, using the template matching procedure and then visually corrected for errors. Data were sampled for at least 5 minutes. Periods of 2-5 minutes were analysed to extract at least 101 events. Where more than 101 events were extracted from a cell, the first 101 events were used for analysis. Number of events used across the neurons were kept constant in order to balance the weight of each neuron in the population data. Data were analysed by comparing average mEPSC amplitudes across cell types, visual deprivation groups, genotype, treatment group (XPro1595) and deprivation timepoints using ANOVA followed by post-hoc t-tests where effects were detected (JMP, SAS software). Linear regression was used to test the strength and statistical significance of correlations. Where data was not normally distributed nonparametric statistics were used. Spine head size data was found to be log-normally distributed, and log transformed before applying parametric statistical methods. Chi-squared tests were used to test the differences in spine-types across cell-types and dendritic branch locations. In those cases where cumulative frequency distribution were plotted, the bin width was determined using the Freedman-Diaconis rule. Matlab, R and Sigma plot software were used for data analysis and plotting graphs.

## Acknowledgments

The authors would like to thank the MRC for funding this research (MR/N003896/1) Frank Sengpiel for critical reading of the text and Stephanie Bagstaff for technical assistance.

