## Supplemental Table 1 for "Hebbian and homeostatic plasticity mechanisms are segregated in sub-types of layer 5 neuron in the visual cortex"

| | No. of Animals | No. of cells – IB | No. of cells – RS | Age range (Days) |
|---|---|---|---|---|
| **Young undeprived** | 30 | 27 | 31 | 27-32 |
| **Old undeprived** | 29 | 27 | 33 | 33-38 |
| **MD 12hr** | 9 | 9 | 9 | 27-30 |
| **MD 3d** | 14 | 8 | 13 | 27-30 |
| **MD 5d** | 10 | 13 | 12 | 33-34 (RS)<br>31-34 (IB) |
| **MD 10d** | 12 | 13 | 14 | 35-40 (RS)<br>34-40 (IB) |
| **DE 12hr** | 11 | 11 | 15 | 27-31 (RS)<br>27-31 (IB) |
| **DE 3d** | 14 | 12 | 14 | 28-31 (RS)<br>28-31 (IB) |
| **DE 5d** | 14 | 10 | 15 | 33-34 (RS)<br>33-34 (IB) |
| **XPro control (12hr & 3d DE)** | 15 | 12 | 17 | 29-33 (RS)<br>28-33 (IB) |
| **XPro 12hr DE** | 11 | 13 | 13 | 28-34 (RS)<br>28-34 (IB) |
| **XPro 3d DE** | 9 | 8 | 13 | 29-32 (RS)<br>29-32 (IB) |
| **XPro control (for 5d DE)** | 9 | 8 | 10 | 31-33 (RS)<br>31-33 (IB) |
| **XPro 5d DE** | 9 | 8 | 11 | 31-33 (RS)<br>31-33 (IB) |
| **T286 control** | 14 | 18 | 16 | 28-34 (RS)<br>28-34 (IB) |
| **T286 12hr DE** | 11 | 11 | 13 | 28-32 (RS)<br>28-31 (IB) |
| **T286 3d DE** | 7 | 12 | 9 | 30-33 (RS)<br>30-33 (IB) |

| | | | | |
|---|---|---|---|---|
| **T286 5d DE** | 11 | 10 | 11 | 32-34 (RS)<br>32-34 (IB) |
| **Spine morphology control** | 11 | 13 | 11 | 27-31 (RS)<br>28-30 (IB) |
| **Spine morphology 12hr DE** | 6 | 10 | 11 | 28-32 (IB)<br>28-32 (RS) |
| **Spine morphology 3d DE** | 7 | 10 | 11 | 28-32 (IB)<br>28-32 (RS) |
| **Total** | 263 | 263 | 302 | |

**Supplementary Table 1.** Number of animals and neurones in each subgroup by age group, genotype, time point, cell type and visual deprivation group (monocular deprivation (MD) or dark exposure (DE)) and treatment group (XPro 1595).
