## Supplementary Figures for "Hebbian and homeostatic plasticity mechanisms are segregated in sub-types of layer 5 neuron in the visual cortex"

### Projection targets

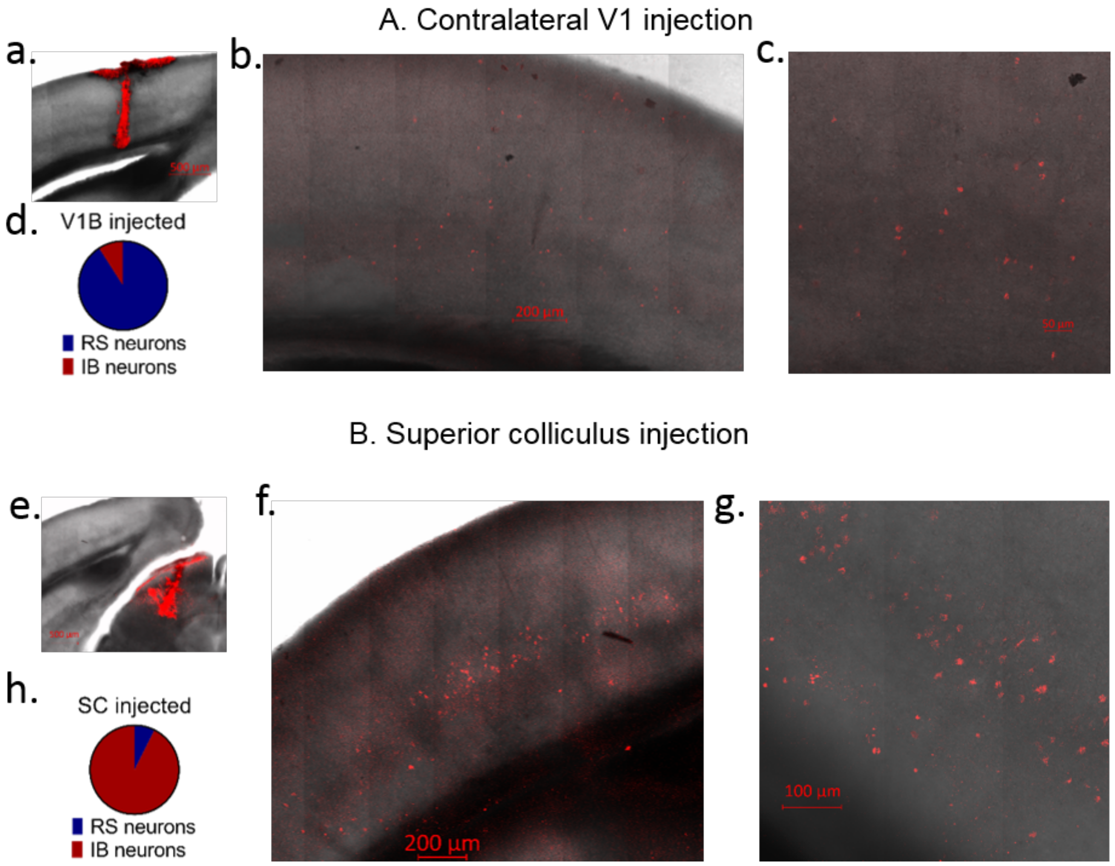


**Figure S1. Coronal sections depicting different projection targets of RS and IB neurons: A.** RS neurons project to contralateral visual cortex, while **B.** IB neurons project to superior colliculus (SC). **a.** Injection of retrobeads in V1b **b.** Labelled neurons in all the layers of visual cortex in the contralateral hemisphere. **c.** Magnified portion of a small area from panel b. **d.** Pie chart representing proportion of RS and IB neurons among neurons projecting to contralateral visual cortex. **e.** Injection of retrobeads in superior colliculus **f.** Labelled neurons mostly present in layer 5 of ipsilateral visual cortex **g.** Magnified portion of a small area from panel f **h.** Pie chart representing proportion of RS and IB neurons among neurons projecting to Superior colliculus.

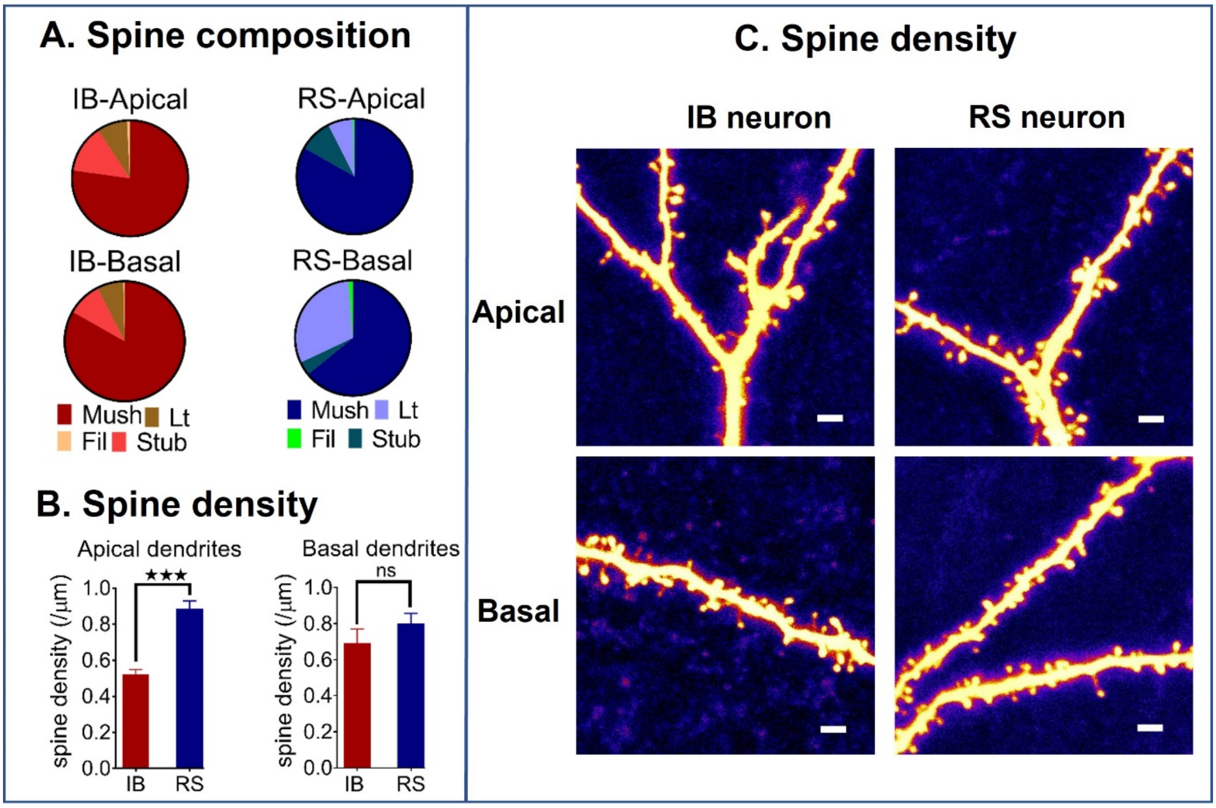


**Figure S2. RS and IB neurons have different spine composition and density: A.** Pie charts showing basal dendrites of RS neurons have more long thin spines than the basal dendrites of IB neurons, while basal dendrites of IB neurons have more mushroom spines than basal dendrites of RS neurons (Mush – Mushroom spines, Lt- Long thin spines, Fil-Filopodia, Stub-Stubby spines). **B.** Apical dendrites of RS neurons have higher spine density than apical dendrites of IB neurons, while there is no different in the spine density of basal dendrites. **C.** Representative images showing the above observations, scale bars – 2 μm.

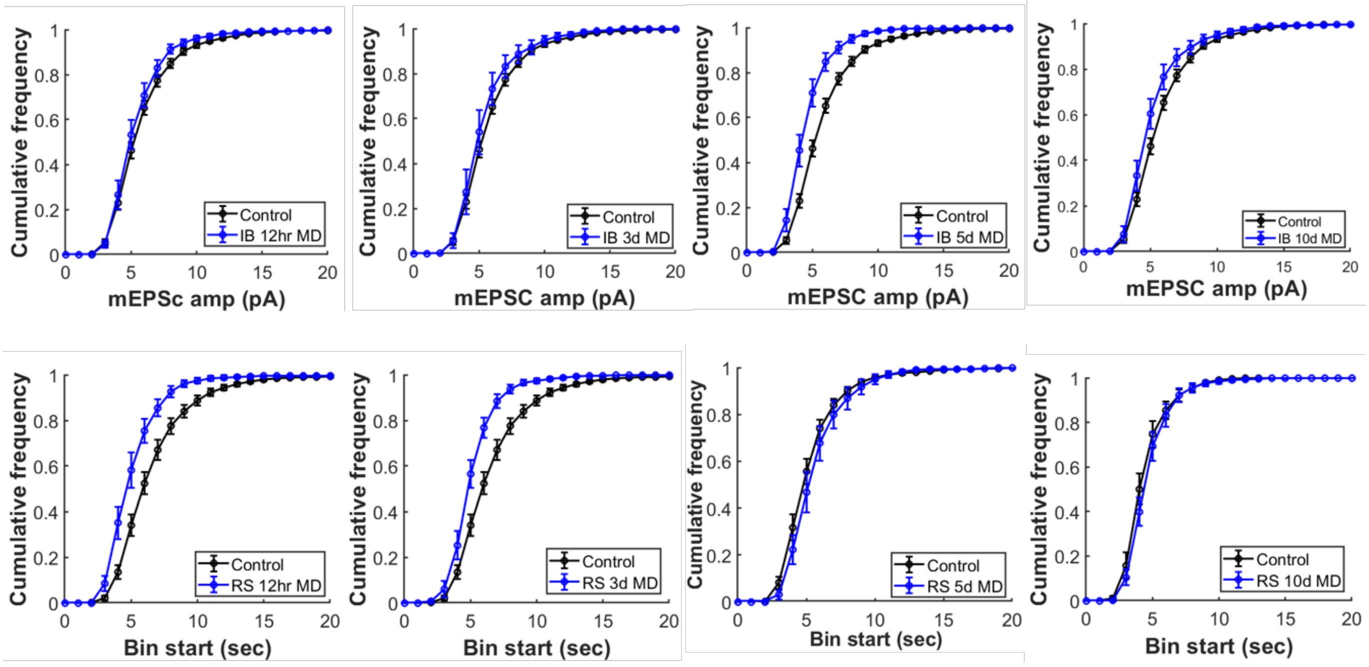


**Figure S3. Cumulative distribution function (cdf) of mEPSC amplitudes in IB and RS neurons with monocular deprivation: Top panel -** cdf of mEPSC amplitudes of IB neurons show reduction in mEPSC amps after 5d MD recovering to baseline after 10d of MD. **Bottom panel -** cdf of mEPSC amplitudes of RS neurons show reduction in mEPSC amplitudes after 12hr and 3d MD recovering to control on 5d MD. Bin interval-1pA.

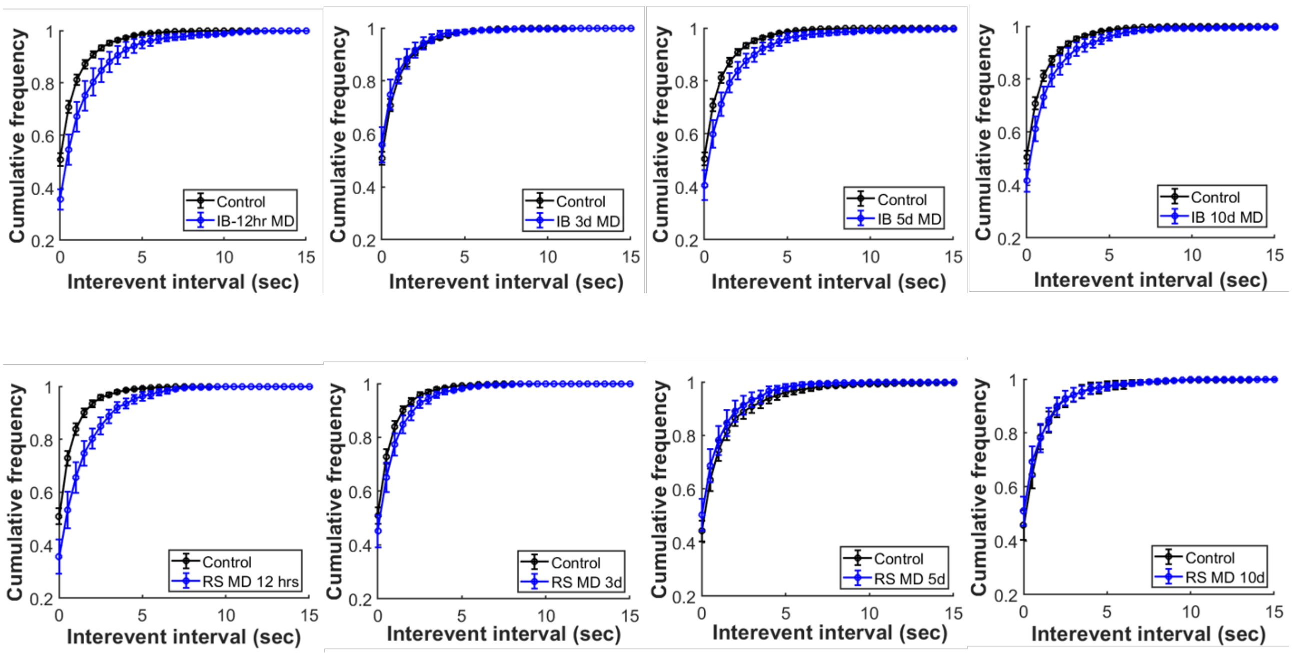


**Figure S4. Cumulative distribution function (cdf) of mEPSC interevent intervals in IB and RS neurons with monocular deprivation(MD): Top panel -** cdf of interevent intervals of mEPSCs in IB neurons does not show any significant effect of MD on mEPSC interevent interval. **Bottom panel -** cdf of mEPSC amplitudes of RS neurons show increase in mEPSC interevent interval after 12hr, and reduction in interevent interval after 5d MD. Bin interval-0.5sec.

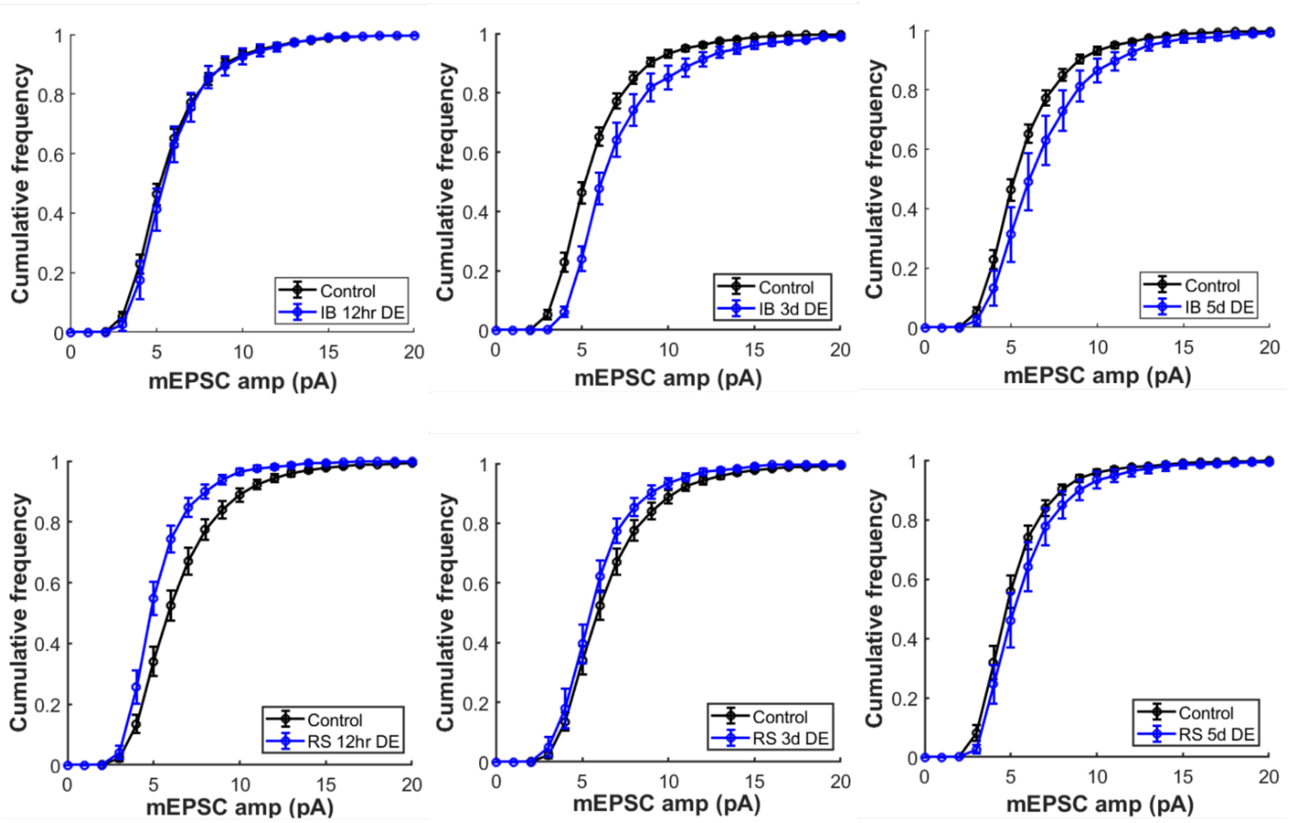


**Figure S5. Cumulative distribution function (cdf) of mEPSC amplitudes in IB and RS neurons with dark exposure(DE): Top panel -** cdf of mEPSC amplitudes of IB neurons show increase in mEPSC amplitudes after 3d DE recovering closer to control data by 5d DE. **Bottom panel -** cdf of mEPSC amplitudes of RS neurons show reduction in mEPSC amplitudes after 12hr of DE and recovering to control values by 3d DE. Bin interval-1pA.

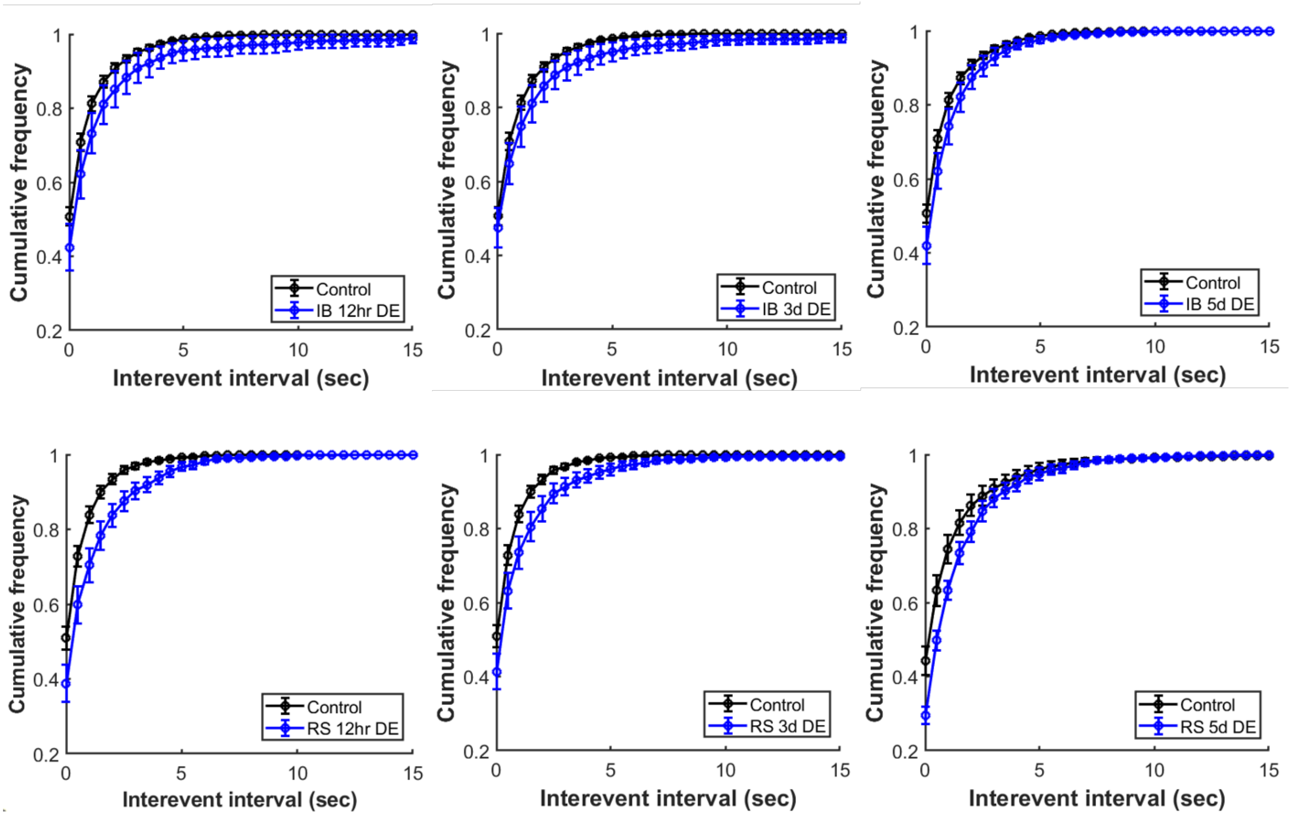


**Figure S6. Cumulative distribution function (cdf) of mEPSC interevent intervals in IB and RS neurons with dark exposure (DE): Top panel -** cdf of interevent intervals of mEPSCs in IB neurons does not show any significant effect of DE on mEPSC interevent interval. **Bottom panel -** cdf of mEPSC amplitudes of RS neurons shows small increase in mEPSC interevent interval at all the time points but the difference is significant only at 5d DE. Bin interval-0.5sec.

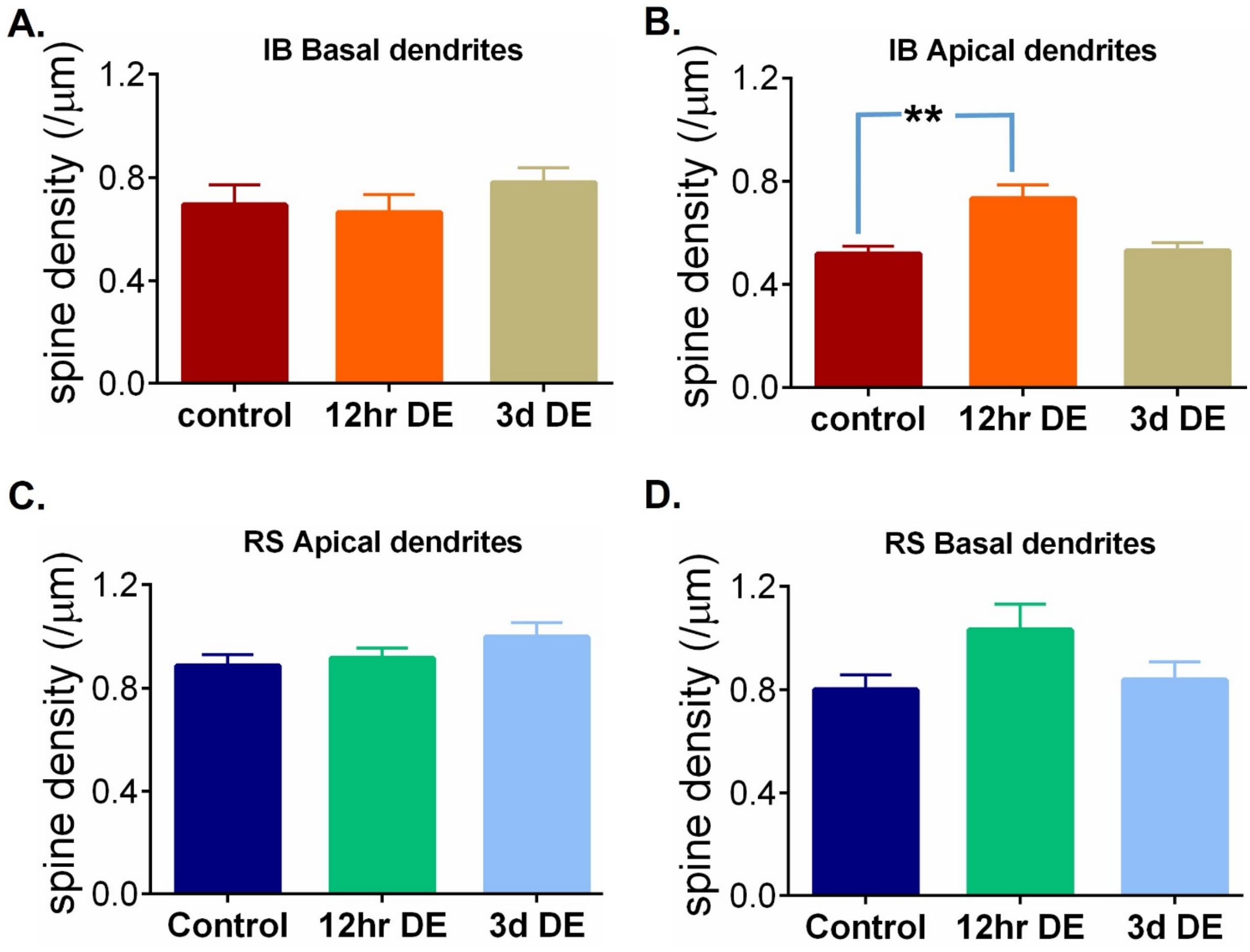


**Figure S7. Effect of dark exposure (DE) on spine density: A.** DE does not change spine density on basal dendrites of IB neurons. **B.** On IB neurons' apical dendrites spine density increases with 12hr DE and comes back to baseline at 3d DE. In RS neurons DE has no significant impact on apical dendrites (**C**), or basal dendrites (**D**).

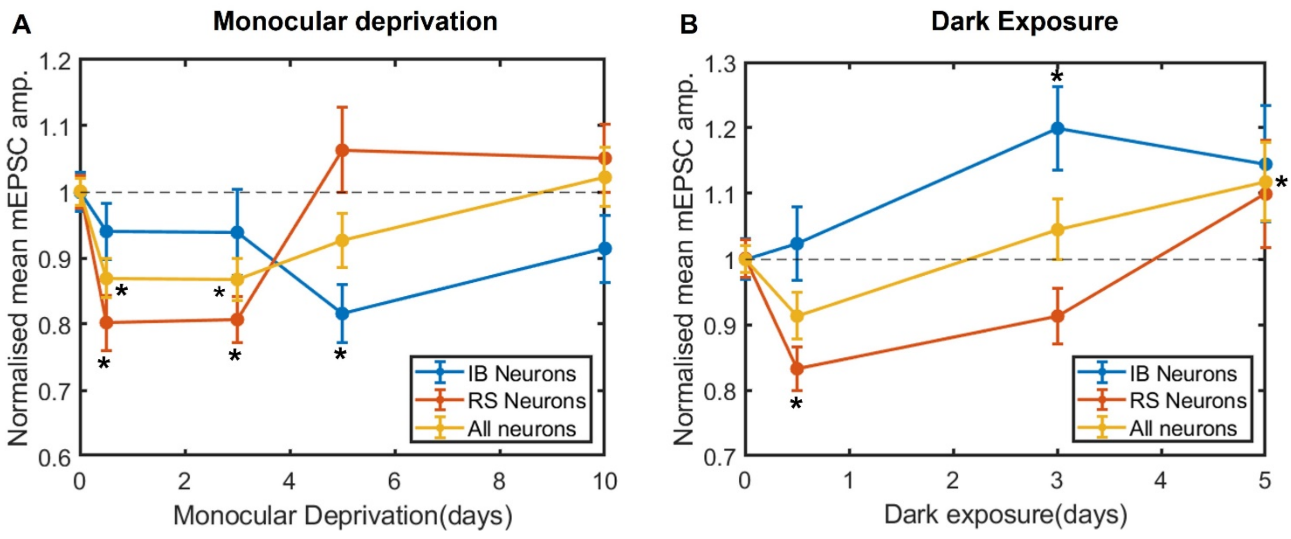


**Figure S8. IB and RS data pooled together for MD and DE: A.** Layer 5 pyramidal neurons show MD induced synaptic depression and slow recovery, if IB and RS neurons were not discriminated (All neurons), while IB and RS neurons' responses to MD are very different than the pooled data and each other. **B.** DE leads to small synaptic depression followed by late potentiation if the IB and RS neurons were not discriminated. Whereas, RS neurons show only depression and recovery, while IB neurons show only potentiation due to DE.
